# Modeling metabolic disease susceptibility and resilience in genetically diverse mice

**DOI:** 10.1101/2025.02.12.637923

**Authors:** Candice N. Baker, Jeffrey M. Harder, Daniel A. Skelly, Isabella Gerdes Gyuricza, Margaret Gaca, Matthew Vincent, Allison Ingalls, Mark P. Keller, Alan D. Attie, Madeleine Braun, Michael Stitzel, Edison T. Liu, Nadia Rosenthal, Gary A. Churchill

## Abstract

Model organisms have provided critical insights into the basic biology of metabolic disorders, however, one of the greatest limitations to translation has been the absence of the genetic heterogeneity that is characteristic of human populations. We examined metabolic health across three genetically diverse mouse strains fed control (low fat, no sucrose) or unhealthy (high fat, high sucrose) diets and observed a wide range of metabolic responses from overt type 2 diabetes in NZO/HlLtJ mice, to obesity and glucose intolerance in C57BL/6J mice, to complete resilience in CAST/EiJ mice. Analysis of multi-tissue gene expression revealed strain- and tissue-specific responses to diet, with strongest responses in white adipose tissue and pancreatic islets. In pancreatic islets, diet response was limited to just NZO mice, which showed high levels of inflammation and associated β cell dysfunction. Adipose tissue was responsive to diet across all three strains and revealed both common and strain-specific changes in inflammatory and metabolic pathways. Using a complementary outbred mouse resource, we mapped genetic loci associated with strain-specific diet responses, including a monocyte regulatory locus on mChr19. This multi-strain approach to modeling metabolic disease revealed a prominent role of white adipose tissue and lipid-associated inflammation in the determination of individual disease risk in response to unhealthy diets.

## Introduction

Metabolic diseases including obesity and type 2 diabetes are endemic in modern human populations. While the rapid increase in prevalence over the past several decades is likely driven by environmental factors including unhealthy diet and sedentary lifestyle, there is a substantial genetic component contributing to individual disease risk^1^. Diet modification and exercise are obvious targets for therapy, but these lifestyle interventions are not always effective, and it is unclear which individuals are most likely to benefit^2,3^. It is also unclear how some individuals avoid weight gain despite having a positive energy balance^4–6^. Studies contrasting obese and lean individuals consuming similar diets have shown that tissue-specific differences in energy expenditure may play a role^7–9^. Additionally, obese individuals exhibit systemic inflammation, which contributes to disease risk^10^. These studies suggest that further investigation of individual variation in response to unhealthy diets may lead to better understanding of heterogeneity in metabolic disease risk and present new opportunities to tailor prevention and intervention therapies to individual needs.

Research using inbred mouse strains has led to critical insights into mechanisms of obesity and insulin resistance^11–13^. Mouse models are well-suited for investigating how environmental challenges such as an obesogenic diet alter the risk of metabolic disease by providing exact control of diet composition and environment, as well as access to tissues to explore molecular and cellular physiology^14–16^. Mouse inbred strains exhibit variability in disease susceptibility and progression, but there has been less focus placed on establishing why some strains develop disease while others display resilience. We propose that systematic characterization of multiple inbred strains to obtain contrasting disease outcomes would provide a disease model that better reflects the genetic heterogeneity of metabolic disease seen in human populations^17,18^.

We carried out comprehensive metabolic phenotyping across three genetically diverse mouse strains, NZO/HlLtJ (NZO), C57BL/6J (B6), and CAST/EiJ (CAST), with distinct responses to high-fat high-sucrose (HFHS) diet. Collectively, these mouse strains constitute a model of heterogenous disease susceptibility. NZO mice are an established model of polygenic obesity and T2D susceptibility^19,20^; they spontaneously develop severe obesity, insulin resistance, and pancreatic β-cell loss^21,22^ in a manner aggravated by harmful dietary factors^23–27^. B6 mice are metabolically healthy on standard diets, but are susceptible to diet-induced obesity, impaired glucose tolerance, and dyslipidemia^28,29^. CAST mice are resilient to diet-induced obesity, insulin resistance, and other features of metabolic disease^15,30^. We profiled six metabolically important tissues using RNA-Seq to identify common and strain-specific transcriptional responses. Finally, we leveraged existing gene expression and clinical trait data from the Diversity Outbred mouse population, a heterogenous mouse stock derived from eight founder strains (including NZO, B6, and CAST) to identify candidate genetic loci that modify HFHS diet response and contribute to individual metabolic disease risk.

## Results

### Development of a heterogeneous metabolic disease model

We fed male and female NZO, B6, and CAST mice either a diet with high-fat and high-sucrose content (HFHS; 44% kcal fat and 1360 kcal sucrose) or a matched control diet with reduced fat content and no sucrose (Control; 10% kcal fat and 0 kcal sucrose) starting at six weeks of age and continuing for nine weeks. To assess the effects of these diets on metabolic health and energy expenditure we carried out comprehensive physiological phenotyping including body weight, food intake, body temperature, body composition, indirect calorimetry, and glucose tolerance (**Fig. 1a**).

**Fig. 1.**
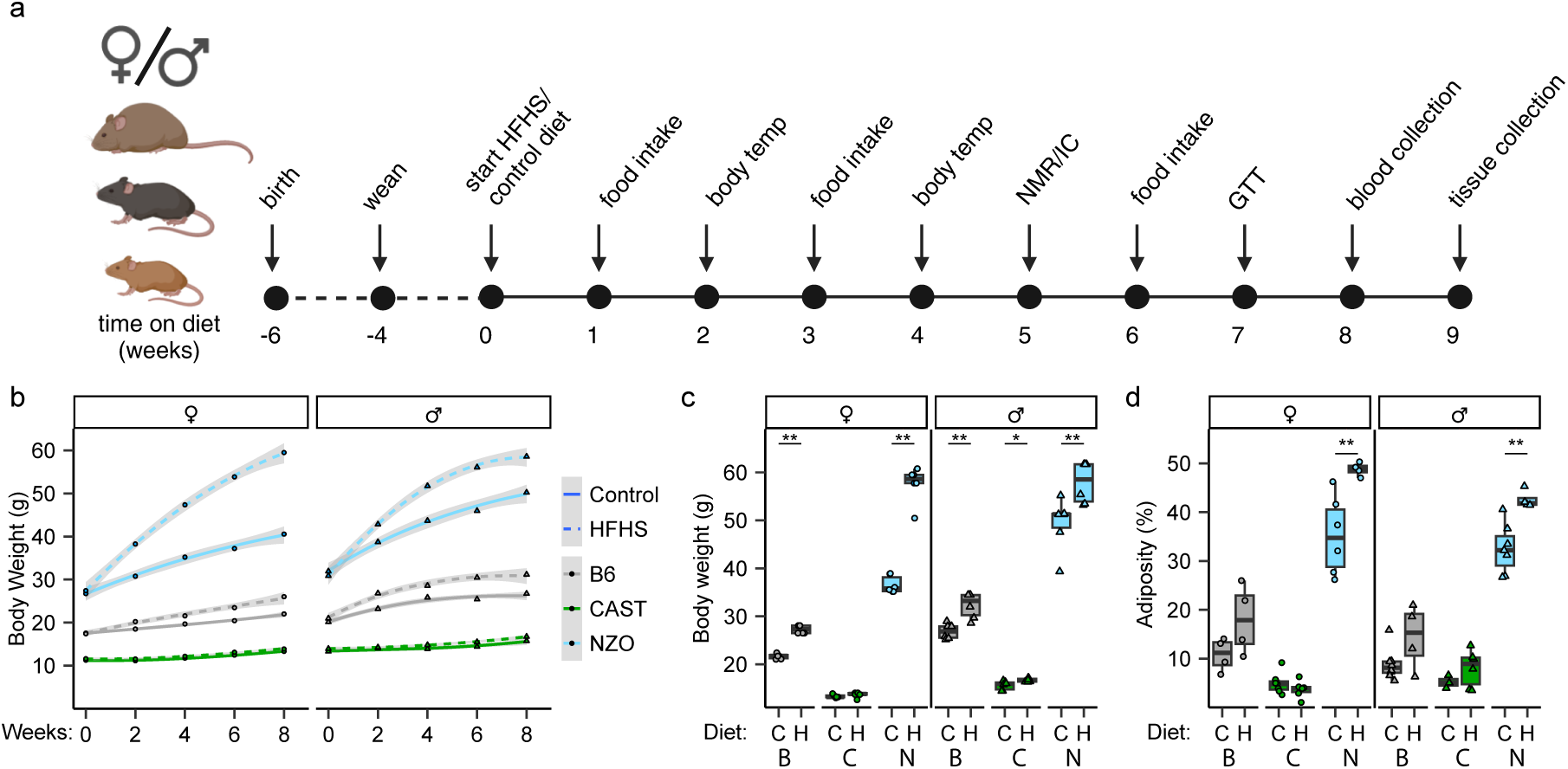
Three genetically diverse mouse strains gained different amounts of body weight and fat when fed a high fat high sugar diet. (**a**) Metabolic phenotyping schedule (n = at least 4 independent mice for each condition, except for glucose tolerance tests n = at least 12 independent mice). (**b**-**c**) Weekly (**b**) and final (**c**) body weight measurements in CAST (green), B6 (grey), and NZO (blue) mice starting at six weeks of age (week 0) continuing for 9 weeks on HFHS, H, (dashed) or control, C, (solid) diets. (**d**) Adiposity (fat mass/total mass x 100) percent after 5 weeks on HFHS (H) or control (C) diet. Box plots show median interquartile range, whiskers indicate 1.5 times interquartile range. Statistical analyses were performed using a two-way ANOVA followed by a Tukey post-hoc test. *P<0.05; **P<0.01. Abbreviations: HFHS or H, high-fat high-sugar; C, control diet; NMR, nuclear magnetic resonance; IC, indirect calorimetry; GTT, glucose tolerance test; B, B6; C, CAST; N, NZO.

#### Diet induced obesity is strain-specific

The HFHS diet led to increased caloric intake in all three strains of mice. Within each strain, mice ate similar amounts of food by weight despite the higher caloric density of the HFHS diet (**Extended Data Fig. 1a-d**). We note that the two diets contained matched amounts of fiber, which may have contributed to similar levels of satiety per gram of food consumed. The HFHS diet promoted weight gain in NZO and B6 mice (**Fig. 1b-c**) but not in CAST mice, as observed in previous studies^15,30^. Lean mass remained unchanged across diets for all three strains, and increases in body weight were due to increased fat mass in NZO and B6 but not in CAST mice (**Extended Data Fig. 1e-f**). After only 5 weeks, NZO mice had increased their already high levels of adiposity (percentage body fat) to 40-50%; B6 mice showed a moderate increase in adiposity; and CAST mice maintained low levels of adiposity on both diets (**Fig. 1d**).

#### CAST mice are resistant to diet-induced glucose intolerance

We measured fasting blood glucose and carried out intraperitoneal glucose tolerance tests (GTT) at seven weeks after the start of diet interventions (**Fig. 2a**). NZO mice on HFHS diet had elevated fasting blood glucose (>175 mg/dL). Most of the male NZO mice and a few female mice displayed overt diabetes (>300 mg/dL fasting blood glucose^19^) (**Fig. 2b**). All NZO mice showed diminished glucose clearance after two hours, and this was exacerbated by the HFHS diet (**Fig. 2c**). Male B6 mice on HFHS diet showed a modest increase in fasting glucose (**Fig. 2b**), and B6 mice of both sexes had a delayed return to baseline consistent with impaired glucose tolerance (**Fig. 2c**). In contrast, CAST mice showed no signs of glucose intolerance across any measure including fasting glucose, 2-hour glucose, and area under GTT curve (AUC) (**Fig. 2a-d**). None of the B6 or CAST mice displayed overt diabetes by either test; however, these two strains had markedly different glucose clearance, which is consistent with HFHS diet-induced glucose intolerance in B6 mice and resistance to glucose intolerance in CAST mice.

**Fig. 2.**
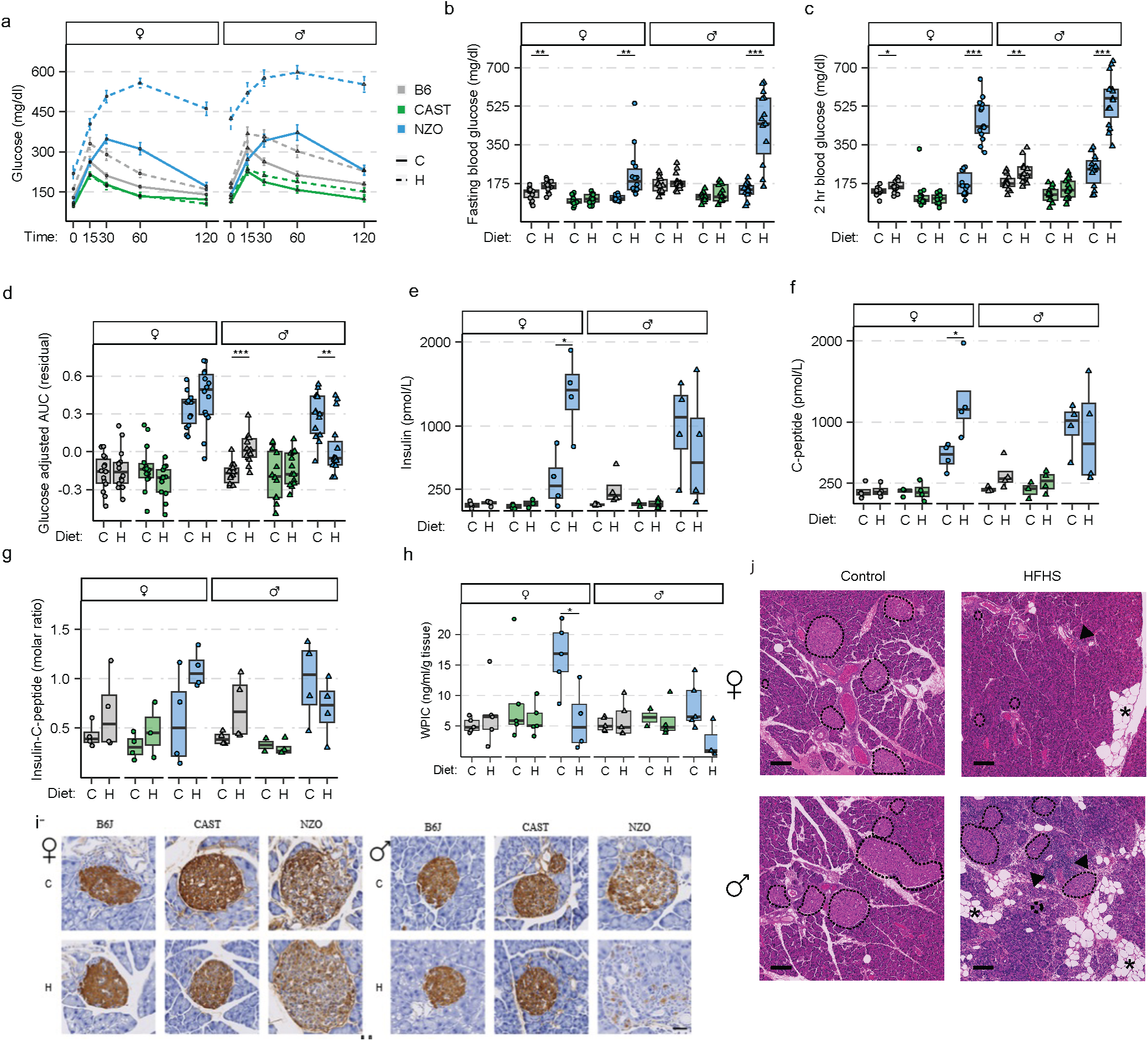
The HFHS diet worsened glucose intolerance and hyperinsulinemia in NZO mice. (**a**) Intraperitoneal glucose tolerance test (GTT) performed after 7 weeks on H or C diet. (**b**) Fasting blood glucose levels after 7 weeks on H or C diet. (**c**) Blood glucose levels two hours after intraperitoneal glucose bolus. (**d**) Adjusted glucose AUC. (**e-g**) Serum insulin (picomole per liter) (**e**) c-peptide (picomole per liter) (**f**) and molar ratio of insulin to c-peptide (**g**) measured after four-hour fast in B6 (grey), CAST (green) or NZO (blue) mice fed HFHS (H) or control (C) diet for eight weeks (n = 4 independent mice for each condition). (**h**) WPIC (nanogram per milliliter per gram of tissue) measured from mice fed H or C diets for nine weeks. (**i**) Representative immunohistochemical stain for insulin on fixed paraffin embedded pancreas samples after nine weeks on H or C diet. Scale bar = 50 µM. (**j**) Representative hematoxylin and eosin staining on NZO mice fed C or H for nine weeks showing islets (dashed circles), lipid droplets (asterisk), and immune infiltration (arrowheads). Scale bar = 100 µM. GTT timepoints are plotted as mean ± s.e.m. Box plots depict interquartile range and median, whiskers indicate 1.5 times interquartile range. Statistical analyses were performed using a two-way ANOVA followed by a Tukey post-hoc test. *P<0.05; **P<0.01; ***P<0.001; ****P<0.0001. Abbreviations: AUC, area under the curve; WPIC, whole pancreas insulin content; C, control diet; H, high-fat high-sugar diet.

#### HFHS diet accelerates pancreatic inflammation and diabetes progression in NZO mice

Hyperinsulinemia is known to arise at a young age in NZO mice and contributes to obesity and insulin resistance^21,22^. We observed elevated serum insulin in NZO mice compared to B6 and CAST after a 4-hour fast (**Fig. 2e**). The HFHS diet increased hyperinsulinemia in NZO females but not in male mice – which may be a consequence of insulin depletion in the more severely affected male NZO mice. NZO mice also had elevated C-peptide levels (**Fig. 2f**) and insulin to C-peptide ratios (**Fig. 2g**), indicating impairment in both insulin release and clearance. In contrast, B6 and CAST mice maintained low insulin levels and insulin to C-peptide ratio regardless of diet. Whole pancreas insulin content (WPIC) was highly variable in NZO mice, consistent with individual variation in disease progression and the timing of β-cell loss (**Fig. 2h**). Reduced insulin content and associated degranulation of pancreatic islets was confirmed by histological staining in NZO males on the HFHS diet (**Fig. 2i**). Using hematoxylin & eosin staining of the pancreas, we observed immune cell infiltration and lipid accumulation within acinar tissue of NZO mice (**Fig. 2j**). Immune cells appeared in highly dense accumulations, either focally or as larger aggregates depending on the mouse. Focal accumulations were primarily observed in NZO females fed HFHS diet or NZO males fed control diet. The largest aggregates of immune cells were within the proximity of lipid droplets in NZO pancreata on HFHS diet. B6 and CAST mice showed no evidence of pathological immune cell infiltration or lipid accumulation in or near islets.

#### Strain differences in energy expenditure were independent of HFHS diet

We next investigated whether differences in HFHS diet-induced metabolic outcomes were linked to strain-specific changes in energy homeostasis. To look at the potential differences in energy expenditure between strains, we performed indirect calorimetry during consecutive periods of *ad libitum* feeding, fasting (24-hours), and post-fasting return to *ad libitum* feeding.

All three strains showed an increased metabolic rate (energy expenditure after regression adjustment for body weight) in response to HFHS diet and differences were largely consistent across feeding states (**Fig. 3a & Extended Data Fig. 2a-c**). We assessed the physical activity of mice by spontaneous running wheel (**Fig. 3b & Extended Data Fig. 2d-f**). NZO mice displayed a strong aversion toward wheel running, reflecting a model of sedentary lifestyle. CAST mice showed the highest level of wheel running, including during the light hours (∼14:00) when mice were normally resting (**Fig. 2b**). We note that CAST mice showed reduced wheel running and lower metabolic rate toward the end of the fasting period on Control diet but not on HFHS diet. Otherwise, mice did not measurably change their overall level or timing of activity between diets (**Extended Data Fig. 2d-f**).

**Fig. 3.**
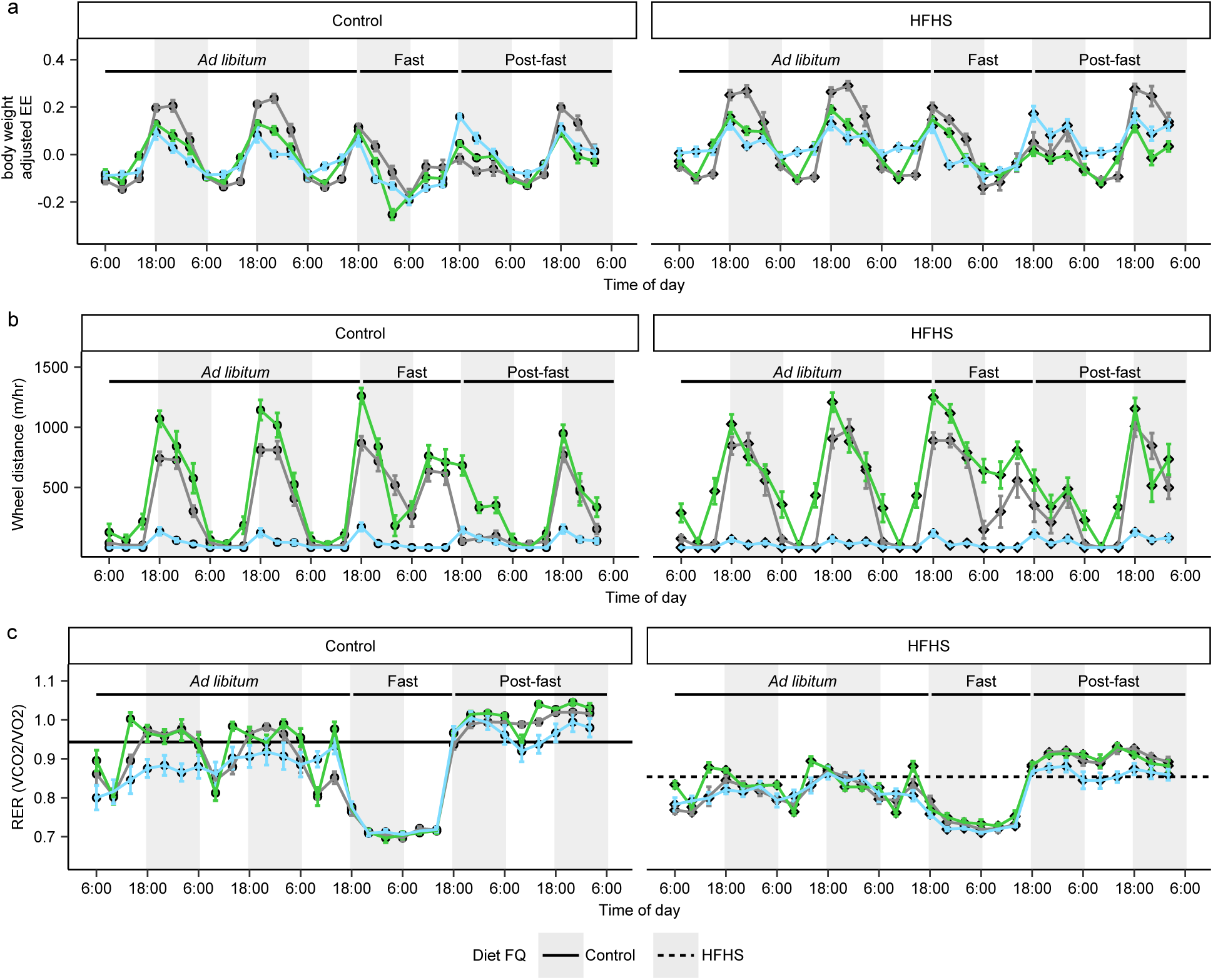
Strain-specific energy expenditure is independent of diet. (**a**) Bodyweight adjusted energy expenditure (metabolic rate) is shown over a 54-hour period with dark/light cycle indicated by shading. Feeding status (top bar) and strain indicated by B6 (grey), CAST (green) and NZO (blue) mice. (**b**) Distance traveled on wheel (meters per hour) over 54-hour period. (**c**) Respiratory exchange ratio (volume of CO2 per volume of O2) over 54-hour period with solid line indicating the food quotient for the control diet and the dotted line indicating the food quotient for the HFHS diet. Error bars indicate means ± s.e.m. Abbreviations: HFHS, high-fat high-sugar; EE, Energy expenditure; RER, respiratory exchange ratio; FQ, food quotient.

The respiratory exchange ratio (RER = the ratio of CO_2_ to O_2_ consumption) obtained from indirect calorimetry provides a measure of energy substrate utilization. RER values near 0.7 indicate the primary energy source is from fatty acids, whereas values of RER near 1.0 indicate utilization of carbohydrates as the primary energy source. Measured RER varied with diet, time of day, and activity level (**Fig. 3c**). We compared the average RER over the *ad libitum* feeding period with energy substrate content of the diets (food quotient (FQ), Control FQ=0.943 and HFHS FQ=0.854; **Extended Data Fig. 2g-i**). On the control diet, NZO and B6 mice were biased toward fatty acid utilization (P=0.002 & P=0.006). Whereas CAST mice aligned their energy utilized with dietary content (P=0.127). Average RER on HFHS diet was reduced as expected and was not significantly different across strains (P=0.15).

All three strains relied on fatty acid metabolism during fasting (with RER ∼ 0.7) and then showed higher levels of carbohydrate utilization upon refeeding (**Fig. 3c** & **Extended Data Fig. 2g-i**). This is expected because fasting lowers insulin levels which can restore insulin sensitivity and carbohydrate utilization during re-feeding. We note that fasting restored carbohydrate metabolism in NZO mice, consistent with improved insulin sensitivity^21^. This effect, however, was short-lived. Within four to eight hours, NZO mice again began to show decreased RER levels indicating a return to fatty acid utilization.

Maintaining body temperature represents a substantial proportion of overall energy expenditure in mice^31^. We observed the lowest body temperature in NZO mice and highest in CAST mice, consistent with observed differences in physical activity and metabolic rate (**Extended Data Fig. 2j**). However, body temperature was not affected by diet in any strain (P=0.28, 0.99, 0.92 in B6, CAST, and NZO mice respectively).

In summary, each strain effectively adapted to the energy substrates provided by HFHS diet without affecting baseline (control diet) differences in energy expenditure. Thus, strain differences in HFHS diet-induced metabolic disease did not appear to be a result of changes in overall energy balance.

### Transcriptional response to HFHS diet is strain- and tissue-specific

We carried out transcriptional profiling of six metabolically important tissues: pancreatic islets, kidney, muscle (gastrocnemius), heart, liver, and white gonadal fat (adipose) (**Fig. 4a**). As expected, Tissue was the main driver of variation in gene expression (**Fig. 4b**). We identified differentially expressed (DE) genes within each tissue using a linear model that included Strain, Diet, Sex, and interaction terms (**Fig. 4c**). Adipose, islet and liver revealed the largest numbers of Diet and Strain x Diet DE transcripts, followed by kidney. Skeletal muscle and heart tissues were least responsive to Diet. To aid interpretation of the Strain x Diet interaction effects, we carried out a second analysis of Diet within each strain to obtain estimates of the strain-specific fold changes and to identify within-strain Diet DE genes (**Fig. 4c**). We looked at overlap of the within-strain Diet DE genes (**Fig. 4d**) and observed that adipose tissue was highly responsive to diet in all three strains with both common and strain-specific transcriptional changes. In contrast, Diet DE genes in pancreatic islets were found only in NZO mice.

**Fig. 4.**
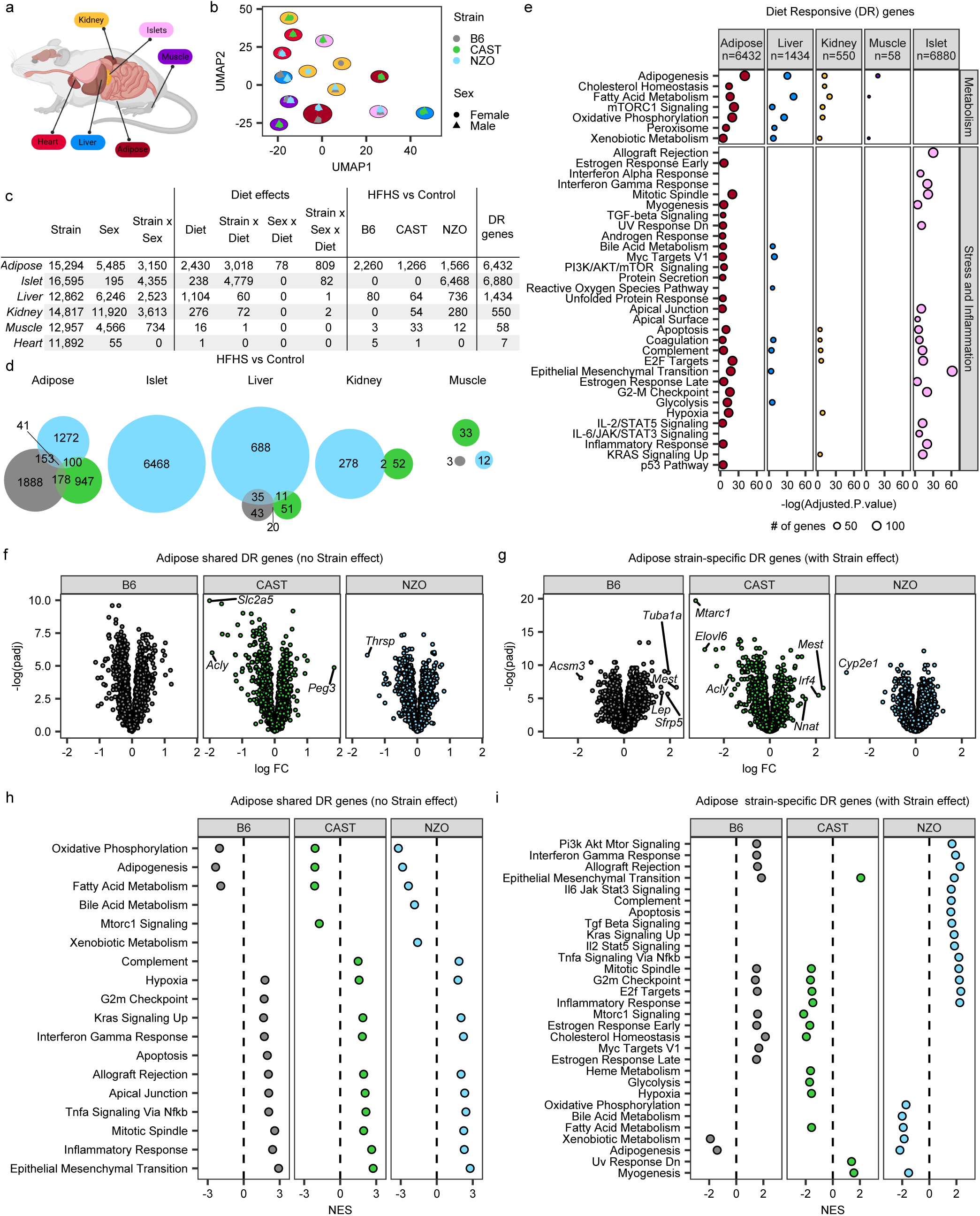
Transcriptional profiling of metabolic tissues reveals genetically driven heterogeneity in the HFHS diet response. (**a**) Tissue collection summary. (**b**) UMAP plot based on the expression levels of 9,114 genes expressed in all tissues and among the top 3,000 most variable genes for at least one tissue. B6, CAST, and NZO mice showed large differences in gene expression in each tissue independent of diet. Colors of the outline ellipses indicate tissues denoted in (**a**). (**c**) Numbers of genes identified by differential gene expression analyses that modeled the effects of strain, sex, diet and their interactions, or compared HFHS vs Control by strain. (**d**) Venn diagrams illustrate uniqueness of the HFHS vs Control diet differential expressed genes for each strain. (**e**) Overrepresentation analyses showed Hallmark MSigDB pathways significantly affected by the diet response. (**f-g**) Volcano plots depic the response of adipose tissue to HFHS diet in B7, CAST, and NZO mice and differences among genes with the largest changes without (**f**) and with (**g**) strain effects. (**h-i**) Enrichment scores derived from GSEA based on diet responsive (no strain effect) (**h**) and strain x diet genes (with strain effect) (**i**) showed that specific Hallmark MSigDB pathways were affected in a common and strain-specific response to HFHS diet.

We compiled a comprehensive list of diet responsive (DR) genes for each tissue based on the union of genes identified in these two analyses and applied gene set enrichment analysis (GSEA) using MSigDB Hallmark pathways (**Fig. 4c**). We identified 38 gene sets that were enriched for DR genes in at least one tissue (FDR<0.05, **Fig. 4e**). Hierarchical clustering of gene sets based on the tissue-specific significance of DR genes revealed a cluster of seven gene sets involved in lipid metabolism that showed a common pattern of Diet response across adipose, liver, kidney, and muscle tissues, but not pancreatic islets. These gene sets reflect changes in fatty acid metabolism and oxidative phosphorylation gene expression consistent with increased lipid oxidation in response to HFHS diet. The remaining 31 gene sets were predominantly associated with adipose or pancreatic islet tissues and involved a range of functions related to cellular stress, immune response, cell proliferation, and glucose metabolism. Next, we looked at diet response within each tissue except the heart, which showed little response to HFHS diet.

<u>*Adipose tissue*</u> had the most complex diet response encompassing 6,432 DR genes enriched across 31 gene sets reflecting diverse functions including energy metabolism, fibrotic remodeling, TGF-β signaling, cellular stress, p53 pathway, unfolded protein response, and a specific type of inflammation – IL2/STAT5 signaling and inflammatory response.

To better understand the common versus strain-specific responses to HFHS diet in adipose tissue, we partitioned the adipose DR genes into those with common effects across strains (n=2,605) (non-significant Strain x Diet interaction) versus strain-specific DR genes (n=3,827) (significant Strain x Diet interaction) and computed gene-set enrichment scores using the fold-change values across each set of genes (common and strain-specific) within each strain (**Fig. 4f-g**). Among the common DR genes, we identified 18 gene sets that were diet responsive across all three strains (**Fig. 4h**), including oxidative phosphorylation, fatty acid metabolism, interferon gamma, TNF-α signaling, and hypoxia gene sets. GSEA of strain-specific DR genes identified 30 gene sets, including most of the gene sets identified separately by GSEA of common DR genes (**Fig. 4i**). A subset of interferon gamma related genes was upregulated in B6 and NZO but not CAST mice. NZO mice had enrichment of more TNF-α signaling genes, along with seven other cytokine and stress response gene sets. Six gene sets were upregulated in B6 and downregulated CAST mice, including Mtorc1 signaling, and cholesterol homeostasis, which have been linked to pro-inflammatory responses by adipose tissue macrophages. Genes involved in glycolysis, which generates both energy and substrates for lipogenesis, were uniquely downregulated in CAST. The fold change analysis suggested that suppressed energy metabolism and elevated inflammatory signaling were common responses to HFHS diet in adipose tissue, while refinement or amplification of these pathways occurred in a strain-specific manner.

In *<u>pancreatic islets</u>*, essentially all 6,880 DR genes identified were specific to NZO (**Extended Data Fig. 3a-b**). We carried out GSEA using fold-change values of all DR genes and found that, despite the lack of multiple testing adjusted statistical significance, there were distinct similarities in the fold-change direction of diet responses between NZO and B6 islets, including 550 shared upregulated genes (log2FC>0.1, unadjusted P<0.0001). GSEA scores for MSigDB Hallmark pathways based on fold changes suggested that functional similarities between B6 and NZO were related to interferon responses. In addition, NZO islets showed a strong upregulation of immune cell signaling, inflammation, and cell death pathways (e.g., IL6, TNFα, IL2, IFN-α, IFNγ, p53, apoptosis) and suppression of pancreatic β-cell pathway genes. These data supported a close relationship between pancreatic inflammation and loss of β-cell identity, a known mechanism of diabetic β-cell failure^32^. Thus, a major effect of the HFHS-diet in NZO mice was the activation of an inflammatory response in islets, which is also present to a lesser degree in B6 mice on HFHS diet In *<u>liver</u>*, 1,434 DR genes were enriched across 12 gene sets (**ED Fig. 3c-d**). We observed a common response to HFHS diet across strains that included increased adipogenesis, fatty acid metabolism, and oxidative phosphorylation. CAST mice showed an upregulation of glycolysis, which could increase glucose usage, helping to regulate blood glucose levels^33^. This contrasts with the CAST response in adipose tissue, where glycolysis was downregulated. In NZO, the liver also showed changes pointing to ectopic fat accumulation and inflammation, such as increased expression of fat storage enzyme *Mogat1* and TNF-α signaling genes^34^.

In *<u>kidney</u>*, 550 DR genes were enriched across 12 gene sets (**ED Fig. 3e-f**). In CAST mice we observed unique upregulation of genes in apoptosis, complement, and coagulation pathways and specific markers of stress or hypoxia, *e.g.*, *Havcr, Igha,* and *Clu*. B6 mice showed an upregulation of inflammatory TNF-α signaling genes, and increased adipogenesis and oxidative phosphorylation that were also seen in NZO mice kidneys.

*<u>Muscle tissue</u>* with 58 DR genes revealed only one gene set enrichment – reduced adipogenesis in CAST mice (**ED Fig 3g-h**). Specifically, in CAST mice, HFHS diet decreased expression of *Acly*, *Elovl6* and *Atp1a3*, whose reduced expression have been associated with a healthier insulin response in people^35,36^.

In summary, these six tissues showed significant differences in transcriptional phenotypes driven by HFHS diet. Metabolic changes were commonly shared between strains but distinct between tissues, consistent with structural differences in energy substrate use. Adipose tissue was the only tissue where HFHS diet caused an upregulation of inflammatory cytokine signaling common to all strains (e.g. inflammatory response, interferon gamma response, TNF-α signaling). In contrast, other inflammatory changes were disease specific (NZO islets – diabetes, NZO adipose tissue – severe obesity). This suggests that the immune response to dietary factors in adipose tissue is an inherent part of dietary adaptation. Strain-specific enriched terms in adipose, liver, kidney, and muscle tissues were mostly shared by B6 and NZO mice but were distinct in CAST mice. Collectively, these data identified several mechanisms that may contribute to metabolic resilience in CAST mice. These mechanisms included regulation of glycolysis in adipose and liver and adipogenesis in muscle, a potential stress or protective response in the kidney, and a unique fine-tuning of the core metabolic and inflammatory responses in adipose tissue (e.g. fatty acid metabolism and interferon gamma response).

### Systemic cytokine levels show limited response to HFHS diet

To better understand the strain differences in systemic inflammation, we profiled serum cytokines (**Fig. 5a**). Levels of obesity-related cytokines such as tumor necrosis factor (TNF), monocyte chemoattractant protein-1 (MCP-1), and interferon gamma-induced protein 10 (IP-10)^37^ were found to be highest in NZO, at intermediate levels in B6, and lowest in circulating CAST serum, independent of diet (**Fig. 5b-d**). Lower serum cytokine levels are consistent with suppression of immune activation in CAST compared to B6 and NZO mice. Notably the levels of these cytokines paralleled expression differences in adipose tissue (**Fig. 5e-g**), rather than islet (**Fig. 5h-j**).

**Fig. 5.**
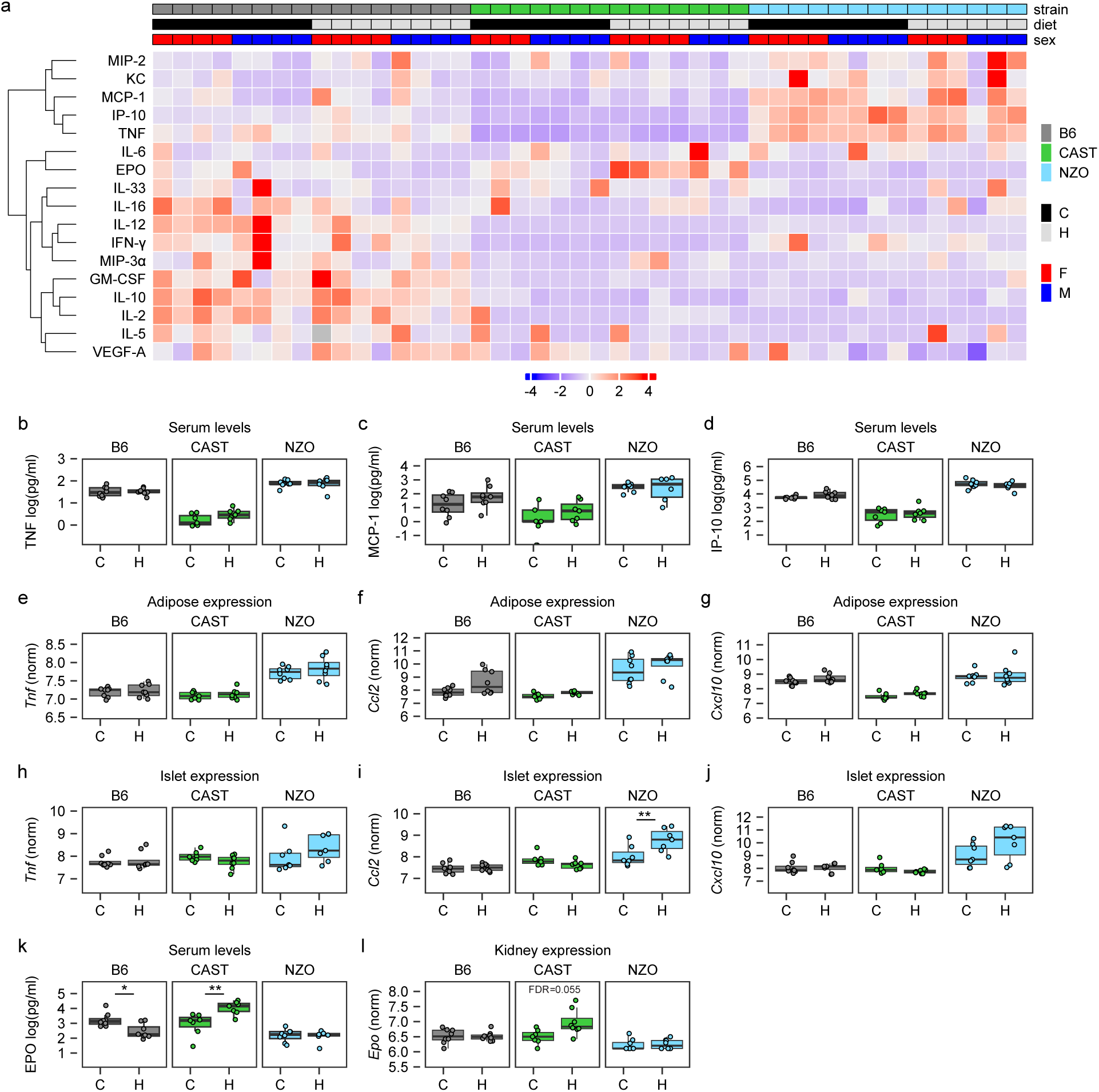
Differential Immune Sensitivity of Adipose Tissue to HFHS Diet Across Mouse Strains. (**a**) Heatmap depicting hierarchical clustering of scaled levels of serum cytokine levels showed large differences between B6, CAST, and NZO on HFHS or Control diets. (**b-d**) Serum levels of TNF, MCP-1 and IP-10 varied between B6, CAST, and NZO mice. (**e-g**) Gene expression of *Tnf*, *Ccl2*, and *Cxcl10* in adipose tissue. (**h-j**) Gene expression of *Tnf*, *Ccl2*, and *Cxcl10* in pancreatic islets. (**k**) Serum levels of EPO in B6, CAST and NZO mice. (**l**) Gene expression of *Epo*, in kidney. Gene expression: **FDR< 0.01; Serum cytokines: *P<0.05, **P<0.01.

Most serum cytokine levels were not directly affected by HFHS diet. However, two did show significant responses to HFHS diet, erythropoietin (EPO; **Fig 5a,k**), which was increased in CAST (P=0.002) and decreased in B6 (P=0.033), and IL16 (**Fig 5a**, P=0.025 decreased in B6). The increased EPO in CAST mice was confirmed by gene expression changes in kidney (**Fig. 5l**).

EPO is a cytokine produced by the kidney crucial for maintaining adequate oxygen levels in the body that also regulates oxidative metabolism in multiple tissues^38^. EPO treatment has been shown to improve glucose tolerance and reduce fat mass accumulation related to obesogenic diets. These findings indicated that HFHS diet related inflammation in NZO adipose and pancreatic islet tissues were tissue-specific processes and that the HFHS diet’s effect in the CAST kidney may contribute to systemic metabolic resilience.

### Strain-specific transcription factor response to HFHS diet in adipose tissue

Adipose tissue had the greatest variety of strain-specific patterns of gene expression changes in response to HFHS diet among the tissues examined. To better understand the drivers of this transcriptional response we examined expression changes in TRANSFAC database annotated transcription factors and counted the number of annotated transcription factor target genes among the set of adipose DR genes. The HFHS diet response was associated with strain-specific expression changes in some of the top ranked transcription factors (**Extended Data Fig. 4a**). In NZO mice, HFHS diet reduced expression of *Elk1* (**Extended Data Fig. 4b**), a transcription factor involved in promoting insulin-mediated adipogenesis^39^. In contrast, CAST increased expression of the interferon regulatory factor 4 with HFHS diet (*Irf4*; **Extended Data Fig. 4c**), which has been linked to increased energy expenditure and improved insulin sensitivity in healthy compared to hypertrophic adipose tissue^40–42^. In B6 adipose tissue, HFHS diet reduced expression of *Xbp1* toward the level observed in NZO (**Extended Data Fig. 4d**), a change associated with increased cellular stress, adipocyte hypertrophy and metabolic disease^43^. The relative expression levels of *Irf4* and *Xbp1* also suggested differences between NZO, B6, and CAST mice in immune cell activation and cytokine production. IRF4 promotes reduced monocyte infiltration and lower cytokine levels in adipose tissue^40^, while loss of XBP1 leads to increased monocyte infiltration and elevated cytokine levels. Thus, overall strain-specific responses to immune activation and cell stress caused by HFHS diet were key factors underlying transcriptional differences.

### Genetic mapping identifies a monocyte regulatory locus on chromosome 19

Based on the findings above, we hypothesized that immune cell infiltration in adipose tissue would vary across strains. Deconvolution of adipose tissue gene expression data using single cell data from Emont *et al*^44^ revealed substantial variation of the monocyte fraction between strains. CAST had an extremely small number of monocytes in adipose tissue, which was unaffected by HFHS diet (**Fig. 6a**). In B6 and NZO mice, the monocyte population was elevated and was further increased by HFHS diet. Together, the strain differences in monocytes, expression of DR transcription factors, and DR gene set enrichment reveal that genetic variation affected both the baseline immune sensitivity and immune changes in response to HFHS diet in a strain-specific manner.

**Fig. 6.**
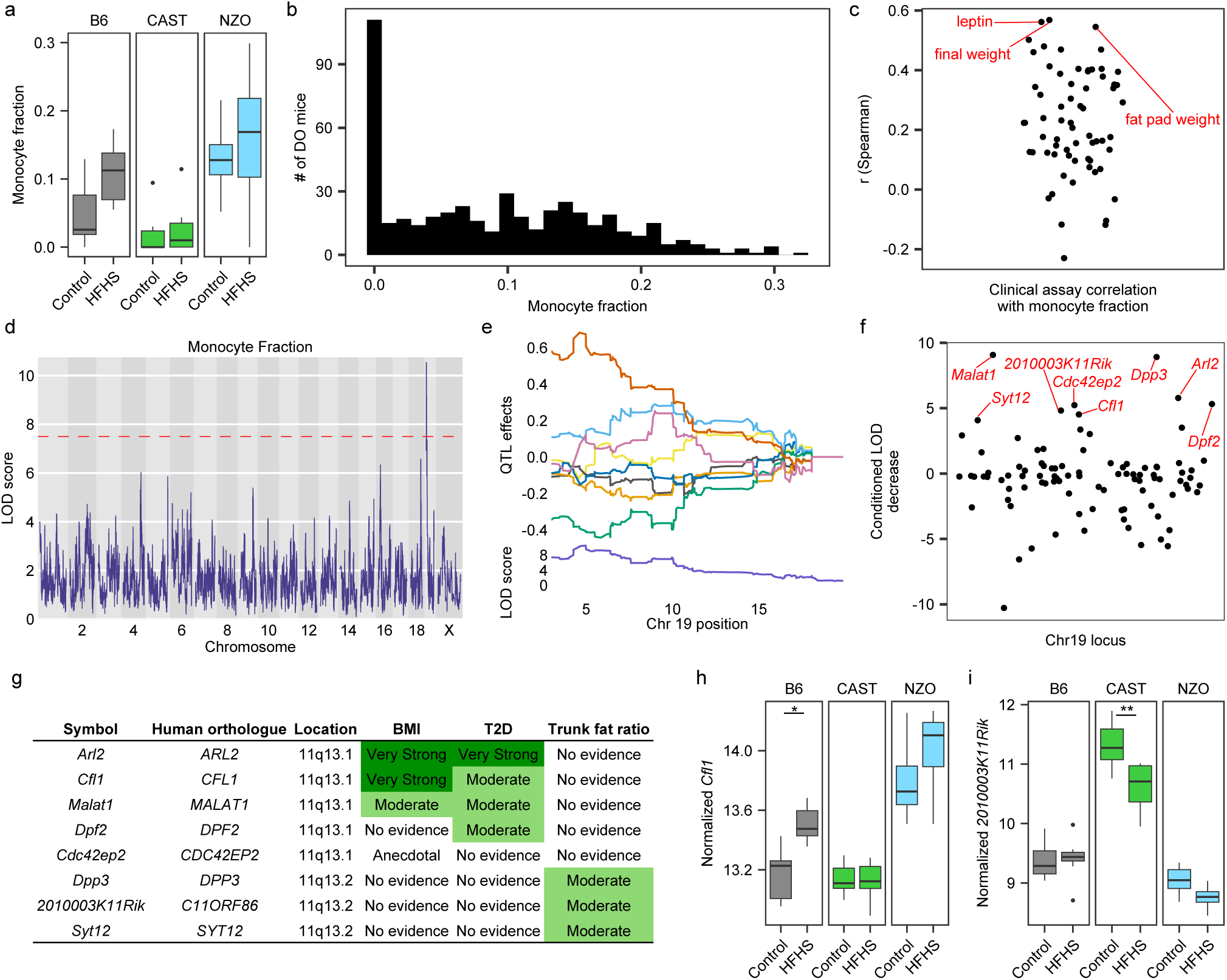
CAST haplotype on Chr19 is associated with reduced monocyte content in DO adipose tissue, shows human-mouse correlation with lower BMI. (**a**) Deconvolution of gene expression data was used to estimate the monocyte fraction in adipose tissue and predicted differences between B6, CAST, and NZO mice. (**b**) The estimated monocyte fraction in DO mice showed variability similar to B6, CAST, and NZO mice. (**c**) Positive correlation of serum leptin, body weight and fat pad weight with monocyte fraction of DO mice. (**d**) LOD scores varied across the genome and a significant genetic association was identified on chromosome 19. (**e**) Allele effects for the QTL on chromosome 19 in A/J (yellow), B6 (grey), 129S1/SvlmJ (pink), NOD/ShiLtJ (dark blue), NZO (light blue), CAST (green), PWK/PhJ (red), and WSB/EiJ (purple) mouse strains. (**f**) Mediation analyses identified eight candidate genes that decreased the LOD score by 4 or more. (**g**) The human region orthologous to the Chr19 locus had very strong genetic evidence for an association with BMI. (**h-i**) Expression of Chr19 locus candidate genes, *Cfl1* (**h**) and *2010003K11Rik* (**i**), *s*howed expression patterns between strains that matched susceptibility to HFHS diet induced obesity. Dashed line indicates LOD=7.4. *FDR<0.05; **FDR< 0.01. Abbreviations: DO, Diversity Outbred; QTL, quantitative trait loci; LOD, logarithm of the odds.

To further investigate the genetic factors that determine this strain-specific immune response to HFHS diet, we obtained RNA sequencing data from adipose tissue (n = 468) of an independent cohort of Diversity Outbred (DO) mice^45–47^. DO mice are an outbred stock derived from eight inbred founder strains that include NZO, B6, and CAST^45,46^. When fed HFHS diet DO mice showed variability in clinical phenotypes related to obesity and diabetes^45,46^.

We applied deconvolution analysis to DO adipose gene expression data to estimate the fraction of monocytes. We identified a subset of DO mice with low monocyte fraction that resembled the CAST phenotype (**Fig. 6b**) and observed that the monocyte fraction was positively correlated with adiposity and body weight across the full cohort of DO mice (**Fig. 6c**). Using the DO mice, we mapped the monocyte fraction to a QTL on chromosome 19 (pos = 4.978 Mb, LOD=10.2; **Fig. 6d**) where the CAST haplotype was associated with a low monocyte fraction (**Fig. 6e**). Mediation analysis^48^ identified eight candidate genes in the locus (**Fig. 6f**). Seven of these candidate genes have human orthologues associated with BMI, trunk fat ratio, or T2D^49,50^ (**Fig. 6g**), consistent with our hypothesis that genetic variation in monocyte responses contributes to metabolic disease susceptibility.

We re-examined the candidate genes from the chromosome 19 QTL in our three inbred strains. We found that *Cfl,* which is a broad mediator of inflammation^51^ implicated in promoting the formation of lipid-associated macrophages^52^, had the lowest expression in CAST and was elevated in response to HFHS diet in B6 mice (**Fig. 6h**). We found that *2010003K11Rik*, or *Faci*, a fasting response gene that has been implicated in fat absorption and promotes leanness^53^, was elevated in CAST adipose compared to B6 and NZO adipose (**Fig. 6i**). We found a decrease in adipose aldose reductase enzyme, *Arl2,* gene expression in B6 mice due to diet (P=0.004, **Extended Data Fig. 6a**). The gene expression changes for synaptic vesicle gene, *Syt12,* were increased in B6 adipose tissue, and conversely, decreased in CAST adipose tissue by HFHS diet (P=0.004 and P=0.02, respectively, **Extended Data Fig. 6b**). Gene expressions for *Dpf2*, *Malat1*, *Dpp3*, and *Cdc42ep2* were unchanged in all strains due to HFHS diet (**Extended Data Fig. 6c-f**). Considering these patterns of strain- and diet-response, as well as their known functions, we propose that one or more of these genes may specifically modulate inflammatory responses to lipids derived from the HFHS diet.

### MHC variation links immune function to metabolic disease risk in mice and humans

To further leverage the DO mice, we looked at variation in adipose tissue expression of strain-specific DR genes. For each mouse we computed an enrichment score for each Hallmark pathway gene set using expression levels of these DR genes (see Methods). We note that many of the pathway enrichment scores were correlated with one another (often between functionally related gene sets) and with clinical phenotypes related to obesity and diabetes in the DO mice (**Fig. 7a**). Most gene set scores were associated with adiposity in DO mice in a manner consistent with how HFHS diet affected NZO, B6, and CAST mice. One notable exception was lower expression of glycolysis or fatty acid metabolism gene sets was associated with greater adiposity in DO mice, but HFHS diet lowered their expression in lean CAST mice (**Fig. 4i**). However, expression of adipogenesis, oxidative phosphorylation, and many cellular stress, and inflammation gene sets correlated with adiposity all aligned with changes seen in NZO, B6, and CAST mice. Genetic mechanisms that regulate these matching expression phenotypes were positioned to contribute to individual differences in adipose tissue expansion and metabolic disease vulnerability across heterogeneous genetic contexts.

**Fig. 7.**
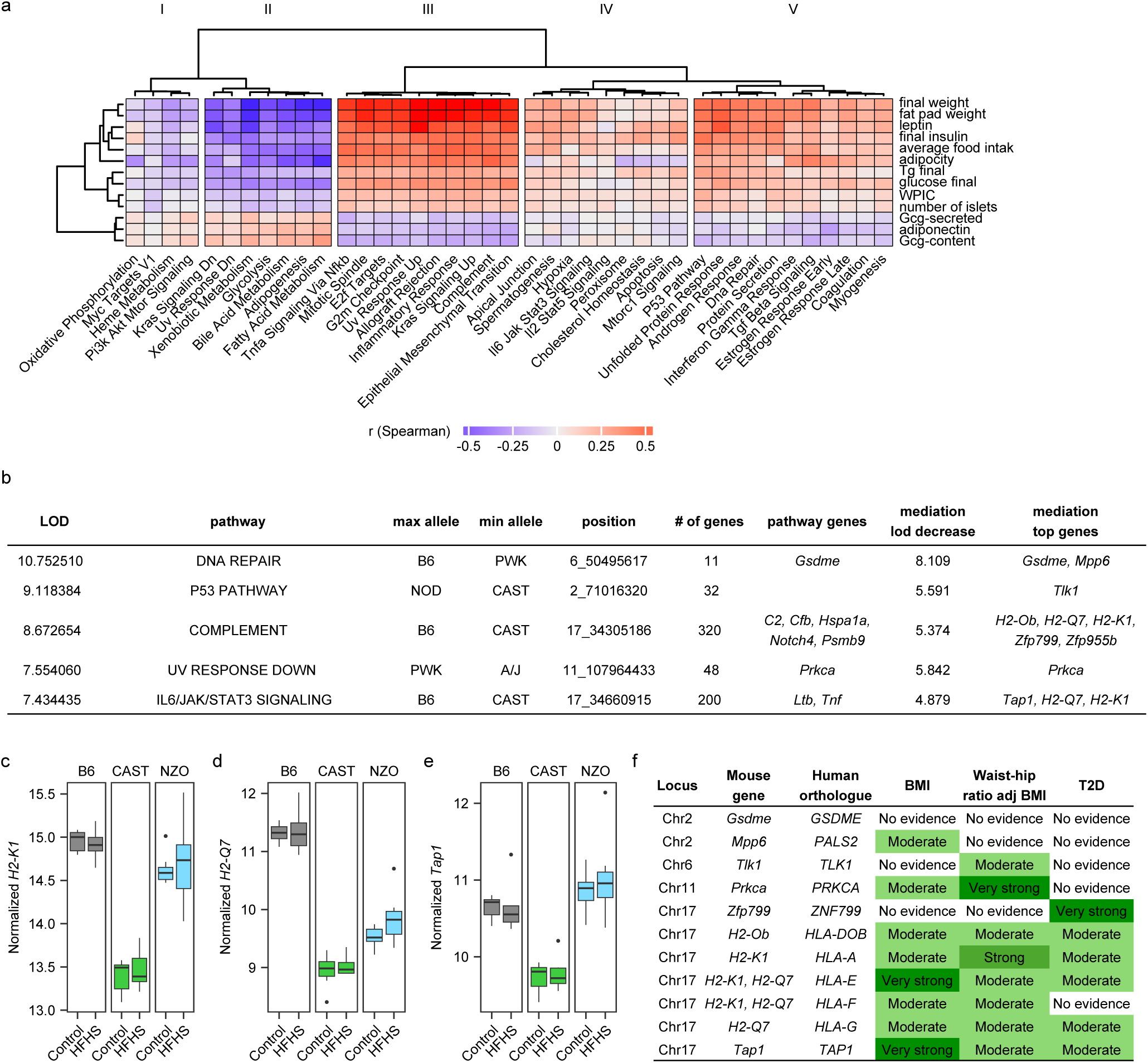
MHC class I gene expression are associated with heightened risk of obesity caused by HFHS diet. (**a**) Heatmap showing correlation between gene set enrichment scores for hallmark pathways using strain x diet genes and various anthropometric and glycemic traits in DO mice. (**b**) Top results of QTL mapping and mediation based on strain x diet gene expression levels in adipose tissue of DO mice fed a HFHS diet. QTL with significant LOD scores were identified for five pathways. Mediation analysis was used to determine whether significance was dependent on the expression of genes within each loci. (**c-e**) Adipose tissue expression of candidate genes in the Chr17 locus, *H2-K1*, *H2-Q7*, and *Tap1*, showed large differences in B6, CAST, and NZO mice, consistent with differences in body weight and obesity. (**f**) Summary of human orthologues and genetic evidence for their role obesity and diabetes related phenotypes. Genetic evidence was determined using the HuGE calculator^49^. Abbreviations: Tg, triglyceride; WPIC, whole pancreas insulin content; Gcg, glucagon.

We identified significant quantitative trait loci (QTL) for five gene set scores and used mediation analysis to identify genes with expression patterns that matched the effects of the QTL (**Fig. 7b** & **Extended Data Fig. 6a-b**). The complement and IL6-JAK-STAT3 signaling pathways shared overlapping QTL and allele effects on chromosome 17 and two major histocompatibility complex (MHC) class I genes, *H2-K1* and *H2-Q7,* were identified as candidate mediators for both loci (**Fig. 7b** & **Extended Data Fig. 6c). The** mouse candidate genes *H2-K7*, *H2-Q1*, and *Tap1,* had elevated expression in B6 compared to CAST mice consistent with allele effects in DO mice (**Fig. 7c-e**). Human genetic evidence scores on the T2D knowledge portal^50^ also implicated MHC class I genes in this locus for association with both BMI and diabetes (**Fig. 7f**). These data indicated that genetic differences in MHC class I gene expression have a major impact on pro-inflammatory responses in adipose tissue, with significant implications for BMI, dietary response, and metabolic health.

Mediation analysis also identified *Gsdme* and *Mpp6* as candidate genes for the DNA repair pathway QTL, *Tlk1* for the p53 pathway QTL, and *Prkca* for the UV response down QTL (**Fig. 7b**). All four loci have orthologous genes that have been linked to body mass index (BMI)^49^ in human genome wide association studies (GWAS). Collectively, the genetic analyses indicate that variation in pathways controlling monocytes, MHC function, and cellular repair and maintenance, contributes to metabolic disease risk across mouse and human populations.

## Discussion

In this study we characterized three genetically diverse mouse strains to reveal heterogeneity in their susceptibility to metabolic disease and differential response to HFHS diet that reflects highly susceptible, intermediate, and resilient genetic backgrounds. By combining physiological and molecular profiling data across multiple tissues, we identified the primary metabolic tissues and biological processes underlying this phenotypic heterogeneity. Our findings highlighted gene expression response in adipose tissue as a primary driver of systemic and tissue-specific inflammation, a key feature of metabolic disease. We then leveraged an independent data resource, gene expression and metabolic phenotyping in outbred DO mice, to genetically map loci that contribute to the heterogeneous disease risk observed in our three strains.

We identified common transcriptional responses across all three strains, primarily in adipose tissue and liver, that match energy use with dietary substrate composition. Adipose tissue showed the strongest and most functionally diverse response to HFHS diet of the tissues examined. We detected 4.5-fold more diet responsive genes in adipose tissue compared to liver and 11-fold more than in skeletal muscle, establishing a hierarchy of diet-responsiveness between metabolic tissues (adipose>liver>muscle>islet). The response to HFHS diet in adipose tissue was larger because of a unique upregulation of genes involved in cellular stress and inflammation, including TNF-α signaling and interferon gamma response. Thus, adipose tissue was the main tissue that showed a conserved immune response to HFHS diet.

Additional diet responses in adipose tissue were strain-specific. HFHS diet affected core response pathways, including TNF-α signaling, interferon gamma, and other stress and inflammation-related pathways in a divergent manner that paralleled strain differences in metabolic disease phenotypes, demonstrating that gene x diet interactions on adipose tissue inflammation play a major role in shaping metabolic health. In an independent population of outbred mice, we confirmed that genetic variation regulated multiple aspects of immune response to HFHS diet, including through regulation of immune cell infiltration in tissues, and recruitment and activation of cellular stress pathways, such as the p53 pathway. Protective CAST haplotypes were associated with low monocyte infiltration and lower levels of MHC-related inflammation in adipose tissue. A drug therapy, PF4178903, that lowers the number of infiltrating monocytes in adipose tissue by blocking MCP1-CCR2 interactions improved metabolic outcomes in B6 mice fed an obesogenic diet^54^. Furthermore, decreasing adipose inflammation in an MCP-1-dependent manner with a T2D drug treatment, pioglitazone, allowed mice to increase calorie intake and not gain weight^55^. Thus, CAST mice appeared to be primed for metabolic resilience by an innate form of immune suppression. The mouse genetic loci implicated here have orthologous human loci and genes implicated in BMI and type 2 diabetes^50^. These data highlight immune sensitivity to dietary factors as a fundamental and modifiable component of body fat accumulation and glucose intolerance that is likely to be evolutionarily conserved and contribute to individual differences. In NZO mice, the immune response was associated with exacerbated diabetes, increasing incidence of prediabetes and diabetes in males and in normally resistant females. The HFHS diet led to an increase in lipid droplet accumulation and immune cell infiltration in pancreatic acinar tissue and downregulation of β-cell identity genes and decreased staining for insulin. Ectopic fat and inflammation are notably recognized as issues in the NZO liver, which has been used to model aspects of non-alcoholic fatty liver disease. Unique TNF-α signaling pathway upregulation was shared by the NZO islets, liver, and adipose tissue. NZO mice also had elevated serum levels of TNF-α compared to B6 and CAST mice. These data suggest NZO mice were highly sensitive to lipid-associated inflammation and TNF-α signaling in a way that contributed to their pathology. Interestingly, many of the obesity candidate genes identified here in adipose tissue inflammation and stress-related QTL have been implicated in both human BMI and T2D. Future research should focus on characterizing the role of these candidate genes as inflammation mediators in type 2 diabetes, particularly by investigating their specific contributions to TNF-α signaling and lipid-associated inflammation in various tissues.

CAST mice displayed response mechanisms related to control of glucose homeostasis that were not present in B6 and NZO mice, including changes in glycolysis regulation in adipose (down regulation) and liver (up regulation) tissues and improved insulin sensitivity in muscle tissue. Based on these findings, the resilience in glucose control seen in CAST was due to either multiple metabolic tissues having unique sensitivity to HFHS diet or coordination by systemic factors. Of note in this regard, CAST mice kidneys showed unique sensitivity to HFHS diet that led to elevated EPO levels. EPO has been shown to improve glucose control in anemic patients^56^ and mouse models^57^ with hyperglycemia. The EPO receptor is expressed in many tissues and cell types and has multiple metabolic and anti-inflammatory functions in other genetic backgrounds (reviewed in [Suresh](https://www.frontiersin.org/journals/physiology/articles/10.3389/fphys.2019.01534/full) *et al*)^38^. In adipose tissue, EPOR signaling inhibits fat accumulation and pro-inflammatory cytokine production by acting on adipocytes and macrophages, respectively. EPO was also suggested to activate JAK2 to promote survival and proliferation of β-cells in the pancreas and to inhibit gluconeogenesis and inflammation caused by a high-fat diet in the liver. Thus, EPO shows potential to act as a general metabolic resilience factor by modifying both metabolic and immune response to HFHS diet. Further work is warranted to establish causality and mechanisms of EPO signaling in response to HFHS diet.

## Conclusion

This study reveals how genetic diversity shapes susceptibility or resilience to metabolic disease, highlighting adipose gene expression and both systemic and tissue-specific inflammation response to HFHS diet as key drivers. Adipose tissue emerges as the central player in diet-induced metabolic adaptation, showing a uniquely robust and diverse transcriptional response—including heightened stress and immune signaling—far surpassing liver, muscle, and islet cells in its role as a determinant of metabolic disease state. NZO mice exhibit extreme sensitivity to lipid-associated inflammation and TNF-α signaling, driving ectopic fat accumulation, immune infiltration, and β-cell dysfunction across multiple tissues. Conversely, CAST mice demonstrated exceptional glucose control linked to unique glycolysis regulation in adipose and liver tissues, enhanced muscle insulin sensitivity, and a novel connection to elevated kidney-derived EPO levels. We identified erythropoietin (EPO) as a potential systemic metabolic resilience factor that coordinates anti-inflammatory and metabolic benefits across tissues—including reducing fat accumulation, protecting β-cells, and inhibiting liver gluconeogenesis—offering a new direction for investigation into therapeutic approaches to combating hyperglycemia and metabolic disease.

## RESOURCE AVAILABILITY

### Lead contact

Requests for further information, resources, and reagents should be directed to Gary Churchill.

### Materials availability

This study did not generate new unique reagents.

### Data and code availability

RNA-seq data have been deposited at GEO and are publicly available as of the date of publication. Accession numbers are adipose; GSE234947, heart; GSE234952, islets; GSE234956 and GSE235481, kidney; GSE234957, liver; GSE235054, skeletal muscle; GSE234960, and DO adipose; GSE266549. RNA-seq data can be visualized at https://churchilllab.jax.org/threebearsalpha/boxplots. Metabolic phenotyping data have been deposited at Figshare and are publicly available as of the date of publication. Microscopy data reported in this paper are shared online using Omero. All original code has been deposited at Github and is publicly available as of the date of publication. Any additional information required to reanalyze the data reported in this paper is available from the lead contact upon request.

## EXPERIMENTAL MODEL AND SUBJECT DETAILS

### Mouse models and *in vivo* experimentation

Mice were maintained and treated in accordance with the guidelines approved by the Association for Assessment and Accreditation of Laboratory Animal Care at The Jackson Laboratory. All animals were supplied by The Jackson Laboratory (C57BL/6J; JR000664, NZO/HlLtJ; JR002105, CAST/EiJ; JR000928) at four weeks of age. Mice were housed in a room free of pathogens at a temperature between 20-22°C and a 12 hr light:dark cycle. NZO and B6J animals were group housed, CAST animals were singly housed due to aggression. Animals were fed either a custom designed HFHS diet (Research Diets D19070208) or control diet (Research Diets D19072203) *ad libitum* starting at six weeks of age. Mice were weighed weekly for the duration of the experiment. Food intake was conducted at 7, 9, and 12 weeks of age by weighing the contents of the grain daily for three days, no differences were observed in the food intake between these three timepoints. Rectal body temperature was measured at 8, 10, and 14 weeks of age by placing the probe of a rodent rectal thermometer (Braintree Scientific) in the rectum of the animal for five seconds daily for three days. No differences were observed in the body temperature between these three timepoints. Nuclear magnetic resonance (NMR) for body composition was performed the manufactures instructions using a 3-in-1 Analyzer (Echo MRI) at 11 weeks of age. A subset of animals at 11 weeks of age were singly housed without enrichment for five days prior to indirect calorimetry testing. Animals were placed in clean cages within the Promethion System (Sable Systems International) with the respective diets for seven days. On the fourth day of testing food was removed for 24-hr then readministered for the remaining 3 days of testing to introduce a fast-refed cycle. Animals had access to voluntary wheel running for the duration of testing. Intraperitoneal glucose tolerance testing (ipGTT) to understand insulin clearance and glucose handling was performed at 13 weeks of age after a 4-6 hour fasting period the baseline measurements were taken by a small tail tip nick using an AlphaTrak2 glucometer and test strips (Zoetis). A bolus intraperitoneal injection of 20% glucose (1g/kg) was administered and repeat tail tip nicks were performed at 15, 30, 60, and 120 min after glucose injection.

### Bulk tissue collection

At 15 weeks of age animals were humanely euthanized by cervical dislocation and tissue (adipose, gastrocnemius, heart, kidney, left liver lobe, dorsal vagal complex, hypothalamus) were harvested and flash frozen in liquid nitrogen for RNA sequencing.

### Mouse islets

At 15 weeks of age animals were humanely euthanized by cervical dislocation and the common bile duct was clamped at the Sphincter of Oddi. A needle was inserted into the bile duct just above the last bifurcation into the liver. Three milliliters of collagenase P (5 units/mL) (Sigma; C7657) and DNaseI (1mg/mL) (Roche; 10104159001) in Hank’s balanced salt solution (HBSS) (Sigma; H1641) were inflated into the pancreas. Then the pancreas was excised and digested in a 37°C water path for 40 min. After shaking the tube for ten seconds ten milliliters of HBSS was added and centrifuged for three min at 180 rcf. The pellet was washed with HBSS twice and then resuspended in 5 milliliters of HBSS. Islets were then handpicked two to three times in a clean petri dish of HBSS. Final pick of clean islets was resuspended in 350 microliters of RLT and 1% β-mercaptoethanol.

## METHOD DETAILS

### Clinical Chemistries

Animals were fasted for four hours prior to serum collection via retro-orbital vein. Whole blood was allowed to sit at room temperature for 30-60 min prior to centrifugation for 5 minutes at 12,500 RPM. Serum was then assayed for glucose (Beckman Coulter; OSR6121), cholesterol (Beckman Coulter; OSR6116), triglyceride (Beckman Coulter; OSR60118), insulin (MSD; K152BZC-1), c-peptide (MSD; K1526JK-1), or 35-plex cytokine panel (MSD; K15083K-1).

### Whole Pancreas Insulin Content

At 16 weeks of age animals were humanely euthanized and the whole pancreas was removed, avoiding any excess fat and mesentery tissue. Pancreas tissue was placed in a pre-weighed 20 mL glass scintillation vial containing acid ethanol (75% HPLC grade EtOH (ThermoFisher; A995-4), 1.5% concentrated hydrochloric acid (ThermoFisher; A144-212) in distilled water). Weight of pancreas is measured for normalization. Using curved scissors, pancreas was chopped for four minutes, and samples were stored at −20°C until all animals were harvested. To obtain insulin measurements, contents of scintillation vials were rinsed with 4 mLs PBS (Roche; 1666789) with 1% BSA (Sigma; A7888), samples were then neutralized with 65 µL 10N NaOH (Fisher; SS255-1) and vortexed for 30 seconds. Samples were then centrifuged at 4°C for 5 min at 2,000 RPM. Samples were diluted 5000X in PBS with 1% BSA and insulin was measured (MSD; K152BZC-1).

### Pancreas Immunohistochemistry

Whole pancreas samples were fixed overnight in 10% NBF then embedded in paraffin for microtome sectioning at 5-micron thickness. Sections were deparaffinized and rehydrated in water. Insulin staining was performed by incubating deparaffinized and rehydrated pancreas sections with mouse monoclonal anti-Insulin + Proinsulin (Abcam; abb8304) at 1:1000 dilution and BOND Polymer Refine Detection (Leica Biosystems; AR9352) for DAB visualization. Aldehyde Fuchsin staining was performed by incubating deparaffinized and rehydrated pancreas sections in a solution of 0.49% Pararosaniline (Sigma; p3750), 1% Paraldehyde (Sigma; p5520), 1.5% 12N HCL (Sigma; 320331) for fifteen minutes. Sections were then counterstained with Mayer’s Hematoxylin (ThermoFisher; 72804) and Eosin (Leica Biosystems; 3801600).

### RNA isolation and QC

RNA was isolated from tissue using the MagMAX mirVana Total RNA Isolation Kit (ThermoFisher; A27828) and the KingFisher Flex purification system (ThermoFisher; 5400610). Frozen tissues were pulverized using a Bessman Tissue Pulverizer (Spectrum Chemical) and homogenized in TRIzol™ Reagent (ThermoFisher; 15596026) using a gentleMACS dissociator (Miltenyi Biotec Inc). After the addition of chloroform to the TRIzol homogenate, the RNA-containing aqueous layer was removed for RNA isolation, according to the manufacturer’s protocol, beginning with the RNA bead binding step. RNA was isolated from pancreatic islets using the RNeasy Micro kit (Qiagen; 74004), according to manufacturers’ protocols, including the optional DNase digest step. RNA was isolated from all other tissues RNA using the RNeasy Mini kit (Qiagen; 74104) and concentrations and quality were assessed using the Nanodrop 8000 spectrophotometer (Thermo Scientific) and the RNA 6000 Pico or RNA ScreenTape assay (Agilent Technologies).

### Library construction

2 ul of diluted (1:1000) ERCC Spike-in Control Mix 1 (ThermoFisher; 4456740) was added to 100ng of each RNA sample prior to library construction. Libraries were constructed using the KAPA mRNA HyperPrep Kit (Roche Sequencing Store; KK8580), according to the manufacturer’s protocol. Briefly, the protocol entails isolation of polyA containing mRNA using oligo-dT magnetic beads, RNA fragmentation, first and second strand cDNA synthesis, ligation of Illumina-specific adapters containing a unique barcode sequence for each library, and PCR amplification. The quality and concentration of the libraries were assessed using the D5000 ScreenTape (Agilent Technologies) and Qubit dsDNA HS Assay (ThermoFisher; Q32851), respectively, according to the manufacturers’ instructions.

### Sequencing

Libraries were sequenced on an Illumina NovaSeq 6000 using the S4 Reagent Kit (Illumina; 20028312). Pancreatic islet libraries were sequenced 150 bp paired-end to a targeted read depth of 100 million read pairs, while all other tissues were sequenced 100 bp paired-end to a targeted read depth of 30 million read pairs.

### Quantification and Statistical Analysis

We used a two-way ANOVA to investigate the effects of HFHS diet on assay measurements across different strains and sexes of mice. The experimental design examined main effects and interaction effects of strain, sex, and diet. Post-hoc analyses were conducted using Tukey’s HSD test to identify group differences for each strain and sex related to HFHS diet while controlling for multiple comparisons. Results were visualized using R packages ggplot2 and ComplexHeatmap^58^.

Differential gene expression (DE) analyses were performed separately for each tissue. Custom R code was written to perform a series of linear model (lm) fits and ANOVA tests to determine the main (additive) effects of strain, sex, and diet, and their interactions. Interaction effects were determined by comparing a full model and a reduced model, excluding one of the interaction terms. To identify DE genes and calculate fold-changes between HFHS and control diet for each strain, multiple comparisons (i.e. B6 HFHS vs B6 control, NZO HFHS vs NZO control, and CAST HFHS vs CAST control) were performed together using the glht function of the multcomp package^59^. P-values for each test were adjusted using the Benjamini-Hochberg method. Results were visualized using R packages ggplot2 and eulerr.

A comprehensive list of diet-responsive genes (DR) was compiled for each tissue using genes with main or interactive ‘diet’ effects in the full model and genes that showed differential expression when comparing HFHS diet and control diet for each strain. An overrepresentation gene set enrichment analysis was performed using a hypergeometric test comparing the DR genes and biological processes defined in the molecular signature database mouse orthologue hallmark gene sets^60^ (MSigDB Hallmark pathways v2020) or regulatory motifs matches from TRANSFAC^61^, accessed in g:Profiler^62^ (version e111_eg58_p18_f463989d). P-values were adjusted using the Benjamini-Hochberg method with a significance threshold of 0.05. Fast pre-ranked gene set enrichment analysis^63^ (fgsea) was performed for each strain and tissue using fold-changes of DR genes and MSigDB Hallmark pathways. Strain comparisons of normalized enrichment scores (NES) were made for pathways with an FDR < 0.05.

For quantitative trait mapping, DO mice were ranked by adipose tissue NES of MSigDB Hallmark pathways, calculated using fgsea of pre-ranked by gene expression values, and by estimating the adipose tissue monocyte fraction with reference-based decomposition using BisqueRNA^64^ and single-cell expression data from Emont *et al*^44^. The NES were based on the 3,018 strain x diet genes (interactive effects) that best captured HFHS-diet associated phenotypic differences in NZO, B6, and CAST mice. Quantitative trait loci (QTL) were found using r/qtl2^65^. The LOD threshold for significant QTL (alpha=0.05) was determined to be 7.4 using 1,000 permutation tests. Mediation analyses to identify candidate genes within the loci were performed based on their expression levels using the R package simicek/intermediate^48^.

## Acknowledgements

Research reported in this publication was supported by The Jackson Laboratory Cube Initiative. We thank Dr. Frederico Rey for advisement on the custom diet. Thank you to the animal caretaker staff at The Jackson Laboratory for their work on these studies. We would also like to thank the contribution of scientific services at The Jackson Laboratory including the Center for Biometric Analysis, Histopathology, Necropsy, and Genomic Technologies for their expert assistance with the work described in this publication.

## Author Contributions

All authors reviewed the final manuscript. Experimentation was performed by CNB, MG, and AI. Analysis and figures were created by JMH, DAS, IGG, MV and GAC. The manuscript was written by CNB, JMH, and GAC. Diversity outbred data was supplied by MK and AA. Conceptualization of project was done by CNB, MB, MS, EL, MS, NAR and GAC.

