## Supplemental Figures for "Modeling metabolic disease susceptibility and resilience in genetically diverse mice"

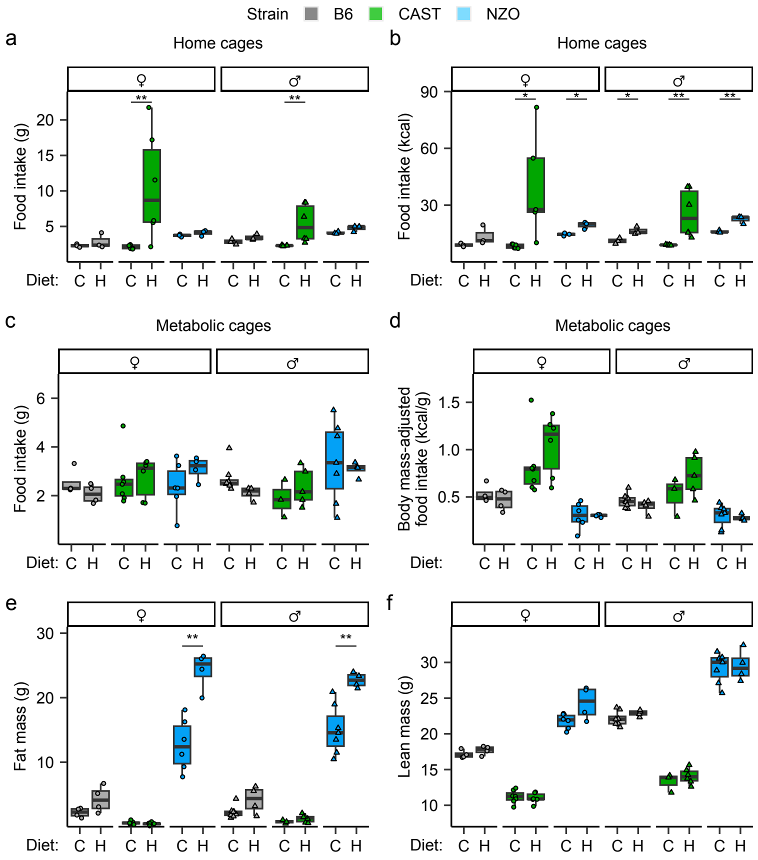


**Extended data figure 1. Food consumption and body composition measurements show strain diversity to HFHS diet.** (**a-b**) Home cage level food intake in grams per mouse (**a**) and kcal per mouse (**b**). (**c-d**) Food intake from metabolic cages in grams per mouse (**c**) and kcal per mouse (**d**). The accuracy of home-cage food intake measurements for CAST mice was compromised by food grinding behavior. However, when eating behavior was monitored in metabolic cages, with singly housed mice, no difference in food consumption between diets was observed (**e**) Fat mass measured by NMR. (**f**). Lean mass measured by nuclear magnetic resonance (NMR). Statistical analyses were performed using a two-way ANOVA followed by a Tukey post-hoc test. *P<0.05 and **P<0.01.

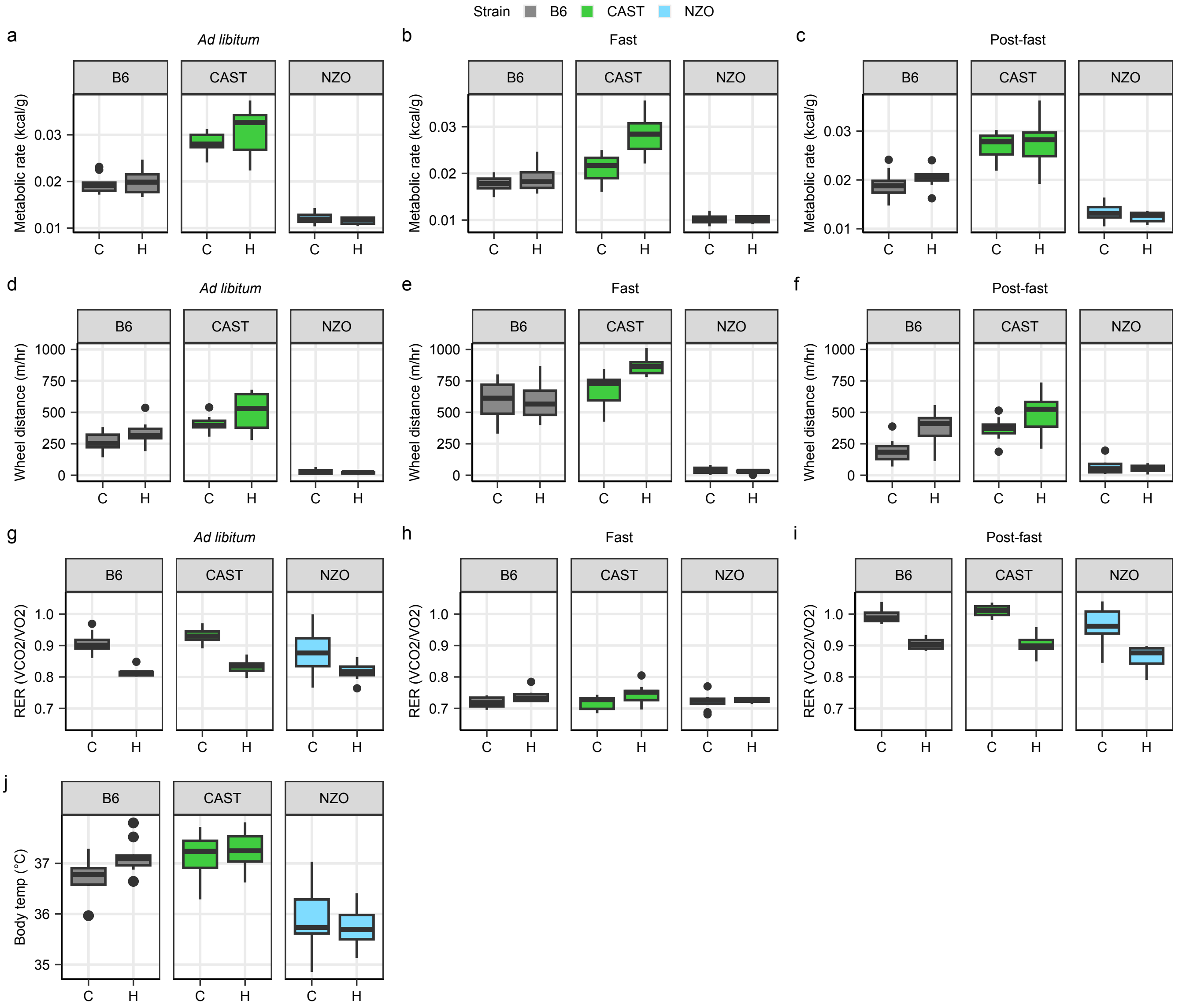


**Extended data figure 2.** (a-c) Metabolic rate (kcal per gram body weight) in ad libitum fed (a), 24 hours fasted (b) and post-fast periods (c) in B6 (grey), CAST (green) and NZO (blue) mice on control (C) or HFHS (H) diets. (d-f) Distance traveled on wheel (meters per hour) in ad libitum fed (d), 24 hours fasted (e) and post-fast periods (f). (g-i) Respiratory exchange ratio (volume of CO2 per volume of O2) in ad libitum fed (g), 24 hours fasted (h) and post-fast periods (i). (j) Average rectal body temperature. Error bars indicate means ± s.e.m. Abbreviations: RER, respiratory exchange ratio; FQ, food quotient.

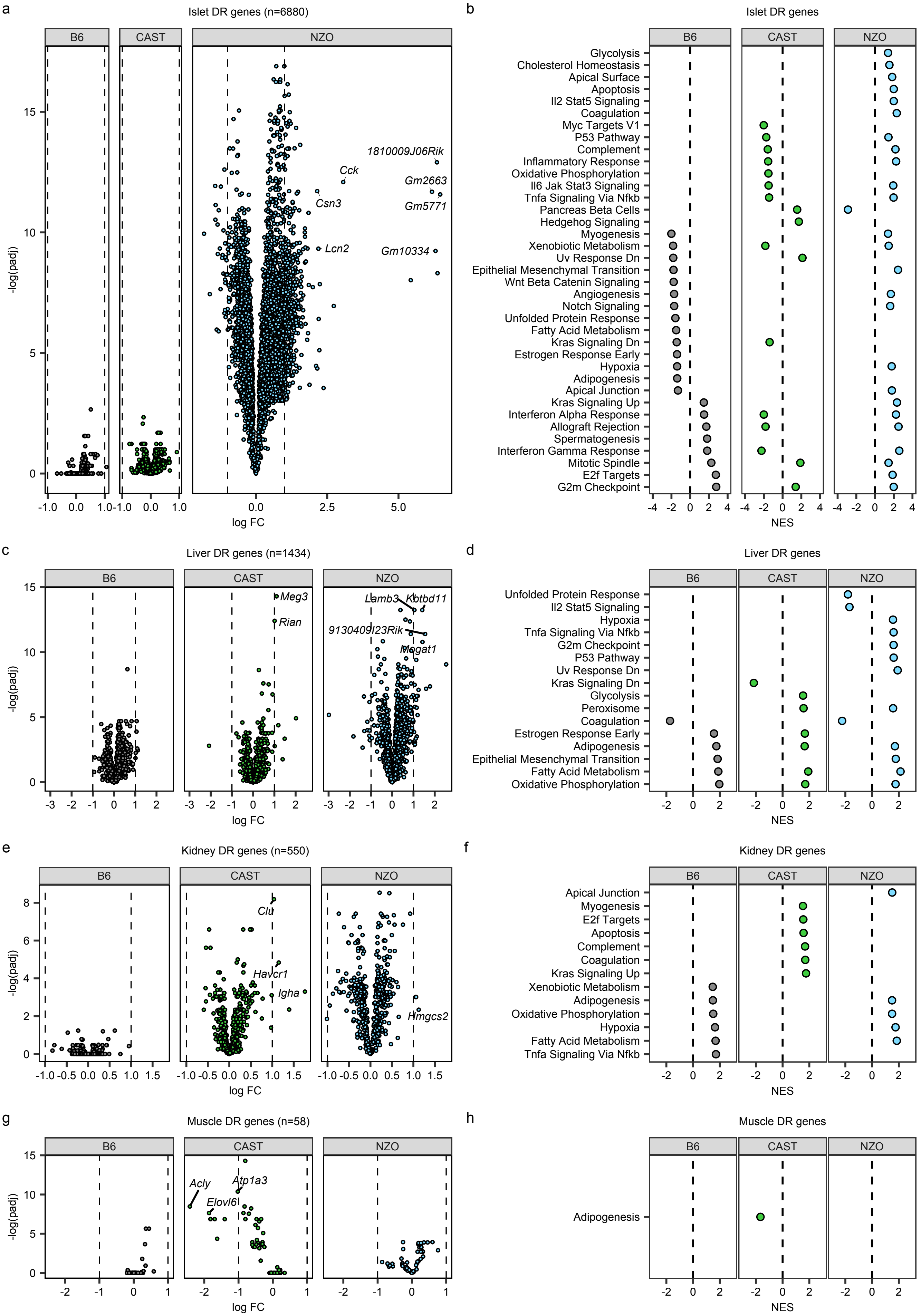
a
Islet DR genes (n=6880)
B6
CAST
NZO
-log(padj)
log FC
1810009J06Rik
Cck
Gm2663
Csn3
Gm5771
Lcn2
Gm10334
b
Islet DR genes
B6
CAST
NZO
Glycolysis
Cholesterol Homeostasis
Apical Surface
Apoptosis
Il2 Stat5 Signaling
Coagulation
Myc Targets V1
P53 Pathway
Complement
Inflammatory Response
Oxidative Phosphorylation
Il6 Jak Stat3 Signaling
Tnfa Signaling Via Nfkb
Pancreas Beta Cells
Hedgehog Signaling
Myogenesis
Xenobiotic Metabolism
Uv Response Dn
Epithelial Mesenchymal Transition
Wnt Beta Catenin Signaling
Angiogenesis
Notch Signaling
Unfolded Protein Response
Fatty Acid Metabolism
Kras Signaling Dn
Estrogen Response Early
Hypoxia
Adipogenesis
Apical Junction
Kras Signaling Up
Interferon Alpha Response
Allograft Rejection
Spermatogenesis
Interferon Gamma Response
Mitotic Spindle
E2f Targets
G2m Checkpoint
NES
c
Liver DR genes (n=1434)
B6
CAST
NZO
-log(padj)
log FC
Meg3
Rian
Lamb3
Kbtbd11
9130409I23Rik
Mogat1
d
Liver DR genes
B6
CAST
NZO
Unfolded Protein Response
Il2 Stat5 Signaling
Hypoxia
Tnfa Signaling Via Nfkb
G2m Checkpoint
P53 Pathway
Uv Response Dn
Kras Signaling Dn
Glycolysis
Peroxisome
Coagulation
Estrogen Response Early
Adipogenesis
Epithelial Mesenchymal Transition
Fatty Acid Metabolism
Oxidative Phosphorylation
NES
e
Kidney DR genes (n=550)
B6
CAST
NZO
-log(padj)
log FC
Clu
Havcr1
Igha
Hmgcs2
f
Kidney DR genes
B6
CAST
NZO
Apical Junction
Myogenesis
E2f Targets
Apoptosis
Complement
Coagulation
Kras Signaling Up
Xenobiotic Metabolism
Adipogenesis
Oxidative Phosphorylation
Hypoxia
Fatty Acid Metabolism
Tnfa Signaling Via Nfkb
NES
g
Muscle DR genes (n=58)
B6
CAST
NZO
-log(padj)
log FC
Acly
Atp1a3
Elovl6
h
Muscle DR genes
B6
CAST
NZO
Adipogenesis
NES

**Extended Data Figure 3. Multi-tissue gene expression analysis reveals tissue specific changes due to diet.** (**a**) Volcano plots showed large differences in the relative magnitude and significance of islet diet response gene expression changes between strains. (**b**) GSEA based on fold changes showed that groups of genes defined by Hallmark MSigDB pathways with largely metabolic and immune related functions were altered in a strain-specific manner in islets by the HFHS diet. (**c**) Volcano plots showed large differences in the relative magnitude and significance of liver diet response gene expression changes between strains. (**d**) Enrichment scores derived from GSEA based on diet response fold changes showed that Hallmark MSigDB pathways were affected in a common or strain-specific manner by the HFHS diet. (**e**) Volcano plots showed large differences in CAST and NZO kidney diet response gene expression changes. (**f**) GSEA based on fold changes in kidney showed groups of genes defined by Hallmark MSigDB pathways. (**g**) Volcanoe plots showing small differences in muscle tissue diet response gene expression changes. (**h**) GSEA based on fold changes in muscle identified adipogenesis as an altered pathway in CAST. Dashed lines indicates logFC=1. Abbreviations: DR, Diet Responsive; FC, fold change; NES, normalized enrichment score.

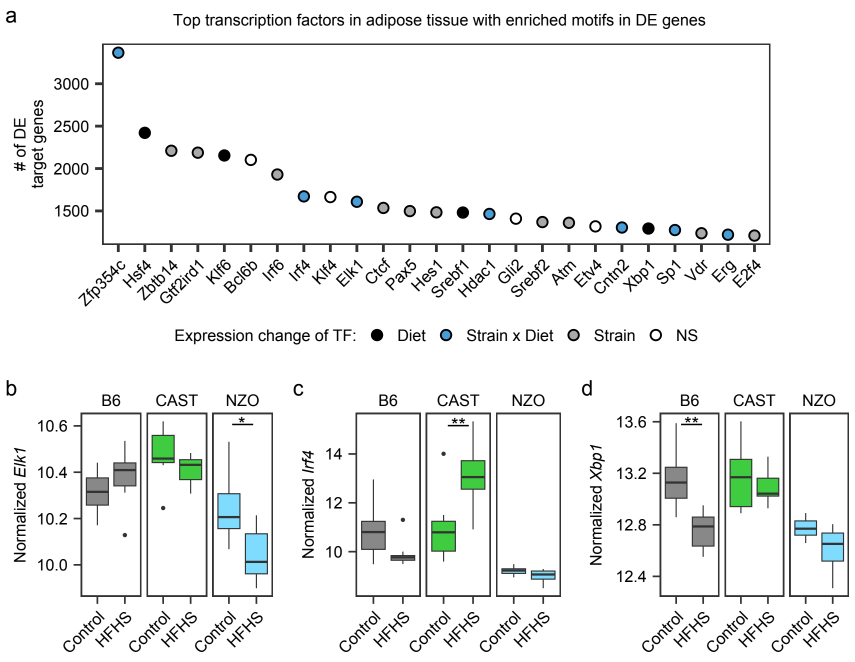


**Extended data figure 4. Adipose tissue transcription factor associated with diet reponse genes.** (**a**) TRANSFAC motif enrichment analysis ranked transcription factors by the number of associated diet response genes in adipose tissue. (**b-d**) HFHS diet and strain differences affected the expression levels of the inflammation and stress response genes *Elk1* (**b**), *Irf4* (**c**) and *Xbp1* (**d**). *FDR<0.05; **FDR<0.01.

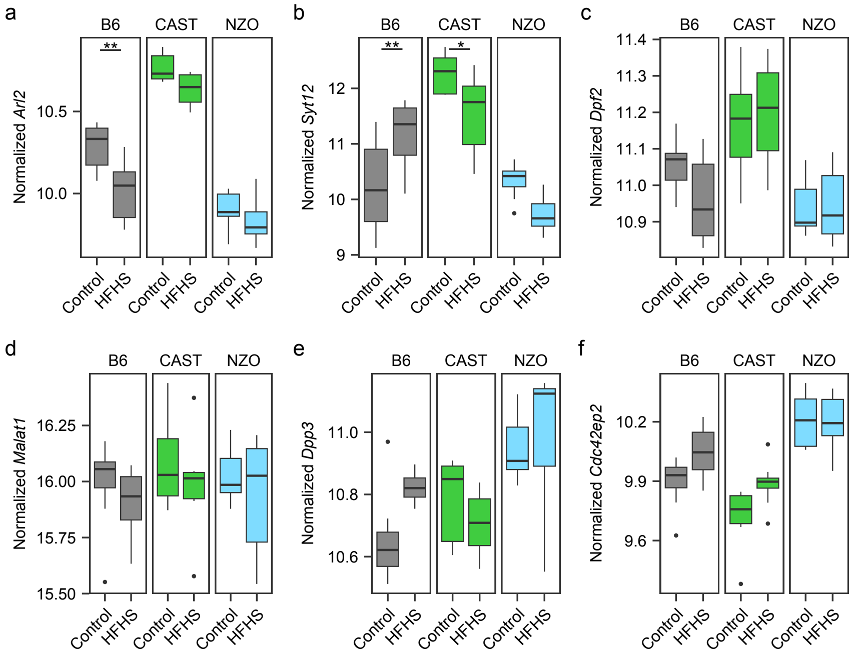


**Extended data figure 5. Candidate genes on Chr. 19 with decreased LOD score.** (**a-f**) Gene expression changes in adipose tissue for *Arl2* (**a**), *Syt12* (**b**), *Dpf2* (**c**), *Malat1* (**d**), *Dpp3* (**e**), and *Cdc42ep2* (**f**) between B6, CAST, and NZO on control or HFHS diet. *FDR<0.05; **FDR<0.01.

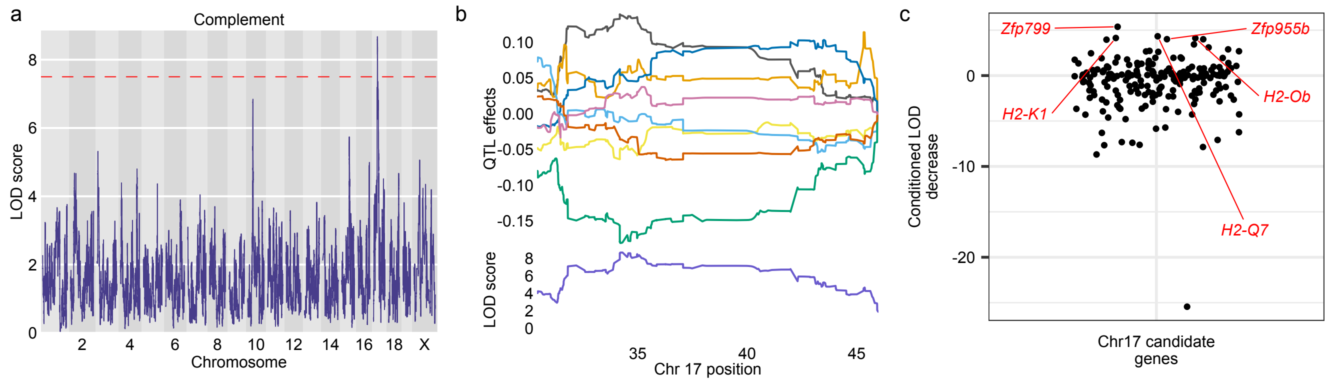


**Extended data figure 6. QTL and mediation analysis for complement pathway in DO adipose tissue.** (**a**) Complement pathway LOD scores varied across the genome and a significant genetic association was identified on chromosome 17. (**b**) Allele effects for complement pathway QTL on chromosome 17 in A/J (yellow), B6 (grey), 129S1/SvlmJ (pink), NOD/ShiLtJ (dark blue), NZO (light blue), CAST (green), PWK/PhJ (red), and WSB/EiJ (purple) mouse strains. (**c**) Mediation analyses identified five candidate genes that decreased the LOD score by 4 or more. Dashed line indicates LOD=7.4.
